# HumanBrainAtlas: an in vivo MRI dataset for detailed segmentations

**DOI:** 10.1101/2022.10.16.511844

**Authors:** Mark M. Schira, Zoey J Isherwood, Mustafa (Steve) Kassem, Markus Barth, Thomas B. Shaw, Michelle M Roberts, George Paxinos

**Affiliations:** School of Psychology, University of Wollongong, Wollongong NSW 2522, Australia; Neuroscience Research Australia and The University of New South Wales, Randwick, NSW 2031 Australia; Department of Psychology, University of Nevada, Reno NV 89557 USA; Centre for Advanced Imaging, The University of Queensland, QLD, 4067 Australia; School of Information Technology and Electrical Engineering, The University of Queensland, QLD, 7067 Australia; School of Psychology, The University of New South Wales, Sydney NSW 2052, Australia

**Keywords:** MRI, human brain, 7T, 3T, T1w, T2w, DWI, image acquisition

## Abstract

We introduce HumanBrainAtlas, an initiative to construct a highly detailed, open-access atlas of the living human brain that combines high-resolution *in vivo* MR imaging and detailed segmentations previously possible only in histological preparations. Here, we present and evaluate the first step of this initiative: a comprehensive dataset of two healthy male volunteers reconstructed to a 0.25 mm^3^ isotropic resolution for T1w, T2w and DWI contrasts. Multiple high-resolution acquisitions were collected for each contrast and each participant, followed by averaging using symmetric group-wise normalisation (Advanced Normalisation Tools). The resulting image quality permits structural parcellations rivalling histology-based atlases, while maintaining the advantages of *in vivo* MRI. For example, components of the thalamus, hypothalamus, and hippocampus - difficult or often impossible to identify using standard MRI protocols, can be identified within the present data. Our data are virtually distortion free, fully 3D, and compatible with existing *in vivo* Neuroimaging analysis tools. The dataset is suitable for teaching and is publicly available via our website (www.hba.neura.edu.au), which also provides data processing scripts. Instead of focusing on coordinates in an averaged brain space, our approach focuses on providing an example segmentation at great detail in the high quality individual brain, this serves as an illustration on what features contrasts and relations can be used to interpret MRI datasets, in research, clinical and education settings.

## Introduction

The investigation of human brain anatomy has a long history, and for most of this time has exclusively relied on the study of post-mortem brains. Today, clinicians and many neuroscientists use Magnetic Resonance Imaging (MRI) to acquire images from living human brains, hence the need to identify brain anatomy in these images. There are a number of projects aiming to provide assistance with brain anatomy such as the BRAIN Initiative (Underwood, 2013), the Human Connectome Project (HCP) (Van Essen et al., 2013), the Big Brain (Amunts et al., 2013; Amunts & Zilles, 2015), the Allen Brain Atlas (Hawrylycz, Lein, Guillozet-Bongaarts, Shen, Ng, Miller, van de Lagemaat, Smith, Ebbert, Riley, Abajian, Beckmann, Bernard, Bertagnolli, Boe, Cartagena, Chakravarty, Chapin, Chong, Dalley, Daly, et al., 2012; Sunkin et al., 2013a), Brainnetome (Jiang, 2013), and the scalable Brain Atlas (Bakker et al., 2015). While these approaches are powerful, there are still major needs faced by contemporary clinicians and researchers not addressed by these projects, principally anatomical resolution. For example, population-bases atlases, by their nature, average out and warp many MR images to a best fit, resulting in loss of detail. As a result, population-based atlases of the human brain typically identify about 50 structures, a handicap for those interested in identifying small structures in an individual brain. Histology based atlases are more comprehensive, identifying as many as 800 structures (J. K. Mai et al., 2016), but histology is limited — by its very nature it is post-mortem and the tissue appearance is significantly different from *in vivo* MR images.

Technological advances have progressively improved the quality and spatial resolution of MRI, including the use of specifically tailored MRI acquisition techniques (MR sequences and protocols), e.g. 3D acquisitions using small, isotropic voxels (Suddarth & Johnson, 1991), which was enabled by the use of higher magnetic field strength (Budde et al., 2014; Pohmann et al., 2016; Ugurbil, 2012) and multi-channel array coils (Roemer et al., 1990)- all measures that improve the signal-to-noise ratio (SNR) dramatically. MRI techniques permit acquisition protocols that provide satisfactory anatomical detail from *in vivo* scans of control subjects, using widely available clinical hardware (Busse et al., 2006; Marques et al., 2010). Advancements now allow levels of resolution that have been previously thought unattainable, recent work demonstrating a high degree of structural information by combining the right acquisition protocols with suitable image processing (Avants et al., 2011; Avants et al., 2010; Janke et al., 2016; Lusebrink et al., 2021). Despite this progress, anatomical delineations of small structures are often inadequate in both cortex and subcortex, with many delineations restricted to gross neuroanatomy, or when being detailed they are often very parochial, covering only segments of the brain (Iglesias et al., 2018; Sone et al., 2016; Winterburn et al., 2013).

While efforts have been made to develop methods for reliable atlas segmentations using minimal, or AI-assisted user-input (Diaz-Pinto et al., 2022; Luo et al., 2021; Zhang et al., 2021) time-intensive manual segmentations on high-quality templates are still considered by many to be the ‘Gold Standard’ for downstream medical imaging processing (Bauer et al., 2013). Atlases not only provide spatial prior probabilities for many segmentation algorithms, but also a reference point for clinical research studies (Ashburner & Friston, 2009; Ashburner et al., 1998; Avants et al., 2010; Awate et al., 2006; Eickhoff et al., 2018; Van Leemput et al., 2003; Wang et al., 2013). When aligning to stereotaxic space, the choice of atlas is often driven by use-case similarities to contrast, field strength, or population characteristics. While there is a push to generate appropriate matching conditional atlases to user input via (for example) machine learning (Balakrishnan et al., 2019; Adrian V Dalca et al., 2019; A. V. Dalca et al., 2019; Hoffmann et al., 2021; Hoopes et al., 2021) or through larger multi-site cohort studies (Fillmore et al., 2015; Richards et al., 2016), there is still a great need for *accurate delineations* of anatomical landmarks in *in vivo* atlases that are of high quality and of similar shape and intensity characteristics as *in vivo* data inputs.

In summary, there is a need for a new, comprehensive, and stereotaxically accurate map of the human brain for *in vivo* neuroimaging applications. The HumanBrainAtlas (HBA) addresses these limitations, combining the anatomical resolution of histology with *in vivo* MRI to remove handicaps each technique possesses. In doing so, it will elevate detail of MRI segmentations to the level of histology. It closes the gap between existing population-based efforts and histology by focusing on the individual instead of the population. Leveraging high quality individual data, also shifts the primary aim. While population based atlases provide a set of coordinates indicating the likely position of structures in some standard space, our approach provides a detailed reference on the anatomical organisation. It provides relative locations and illustrates the contrasts in *in vivo* MR images, it links postmortem techniques, to in vivo MRI.

To this end, HBA renders two living subjects in ultra-high-resolution MRI of 250 microns. As in the histological atlas of Mai, Majtanik and Paxinos (J. K. Mai et al., 2016), we aim to define approximately 800 structures, providing similar accuracy for science and clinical practice, but within the much more ubiquitous and clinically relevant space of *in vivo* MRI. Our ambition is to link the field of postmortem anatomy to *in vivo* MRI. For this goal, we present herein high-resolution MRI data at 7T (T1w and T2w) and 3T (DWI). The general approach was to collect repeated images at maximum permissible resolution and average these individual, grainy images at even greater resolution to construct a super resolution average of individual brains, bringing the neuroanatomical resolution of histology to the world of MRI. The datasets, post-processing protocols and ongoing progress of delineations are made available for open access through our website hba.neura.edu.au.

## Results

Averaging multiple acquisitions through the ANTs multivariate template fitting resulted in significant quality improvement — from grainy high-noise single acquisition images to an excellent average within an individual subject (Figure 1).

**Figure 1.**
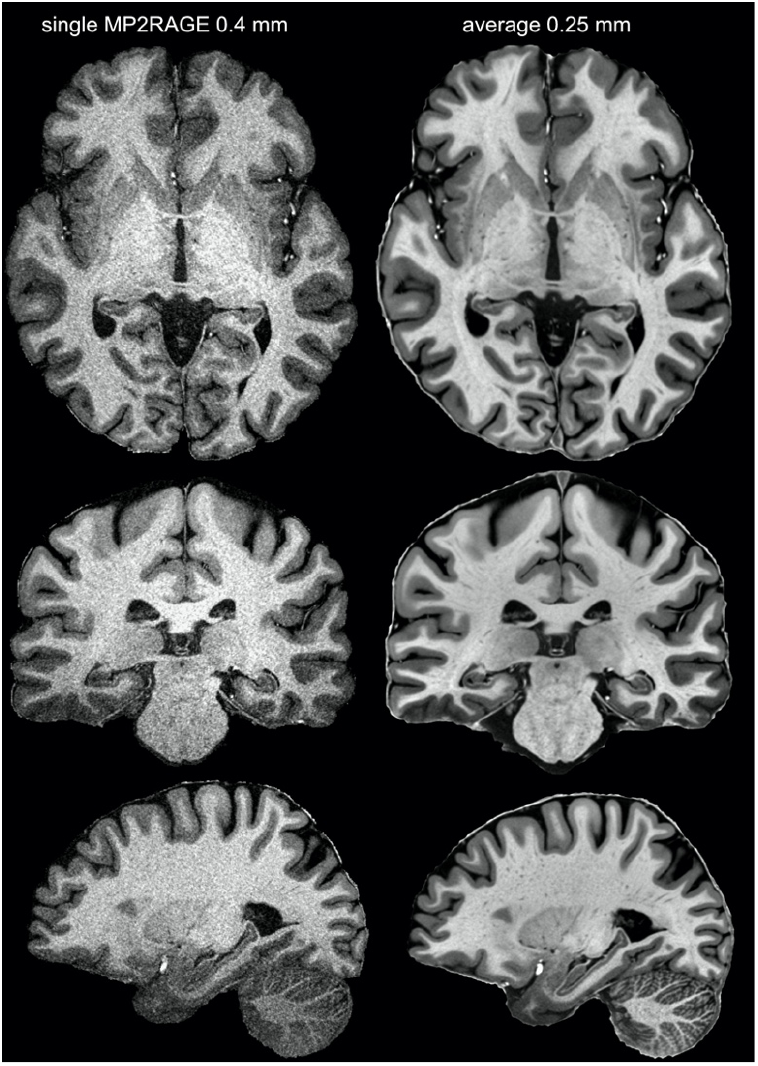
From single acquisition to high-resolution and quality average. The left panel shows the denoised and skull stripped (UNIDEN) image of a single MP2RAGE acquisition, while the right shows the average T1w image in 0.25mm resolution. The left image shows considerable detail; this detail is masked by strong grain (noise). On the other hand the images on the right are virtually noise free and even small contrast variations can be relied on, revealing meaningful structural detail.

Our results also demonstrate fine detail in our final images, detail unavailable in the initial scans. They also highlight the importance of high-resolution for imaging structural details, for example in the hippocampus (Figure 2). Attempting to discern hippocampal subfields at 1 mm involves risky guessing (Wisse et al., 2021) even in excellent data as illustrated in Figure 2; whereas, at 0.25 mm^3^ resolution subfields are clearly discernible in T1w images.

**Figure 2.**
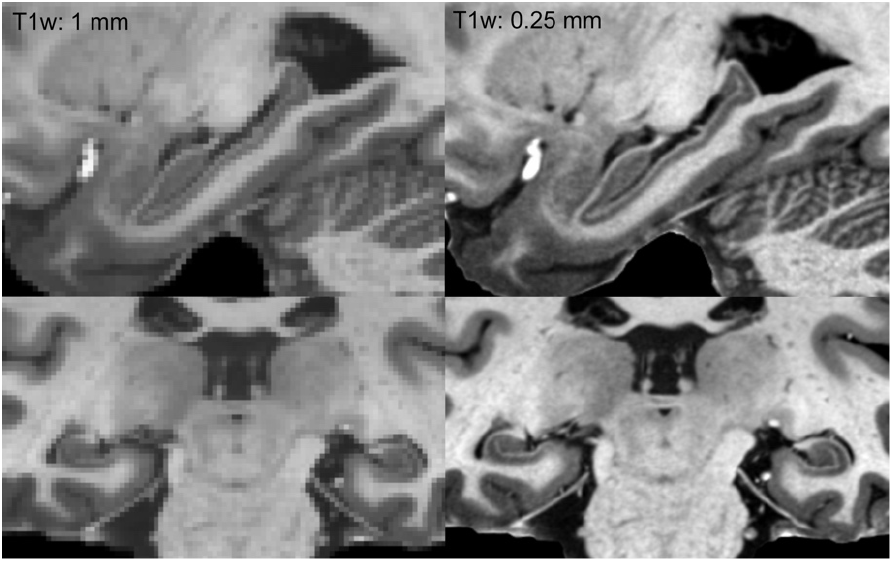
The importance of resolution in segmenting the hippocampus. The left column shows the dataset at 1 mm resolution, while the right column shows the exact same data at its original resolution of 0.25 mm.

Our results also demonstrate the ability of post processing to reveal structural detail finer than the acquisition resolution. Figure 3 shows the resolution gain achieved by the super resolved processing for the FAC data. DWI was acquired at 1.25 mm isotropic; while this is a high-resolution for DWI, it is low compared to our T1w and T2w protocols. Comparing the acquisition resolution (bottom right) and with the high-resolution average in 0.5 mm^3^ (bottom left) demonstrates the detail that was achieved through our post-processing.

**Figure 3.**
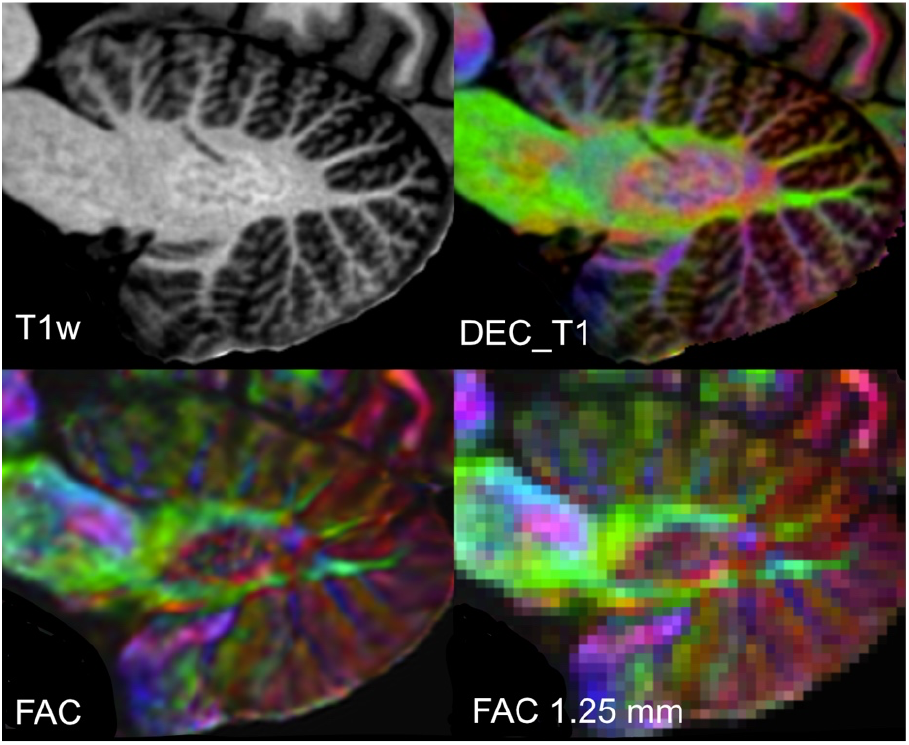
Sagittal plane showing the cerebellar dentate nucleus across a selection of contrasts. The dentate can be clearly seen in both the T1w and FAC example, but most clearly in DEC_T1 which combines these two image modalities. Lastly the bottom right panel shows the FAC at 1.25mm resolution. This demonstrates the superior detail revealed in the left FAC image. (Red: LR axis or vice versa, blue IS green AP, the same colour coding applies to all FAC or DEC_T1 images).

Having demonstrated that our analysis pipeline improves the available resolution, the next relevant question is whether the structural information revealed is sufficient to support accurate and comprehensive delineations of brain structures. Figure 5 focuses on the axial plane through the AC-PC line (z=0) on which we delineate 52 structures, for example revealing substructures of the globus pallidus and its flanking structures the putamen and the internal capsule. Visible are the thalamic substructures such as the ventral anterior nucleus, lateral thalamic nuclei, the medial geniculate nucleus and the pulvinar. Posterior to the thalamus can be seen the hippocampal subregions — dentate gyrus, CA1, subiculum, and pre- and parasubiculum.

**Figure 4.**
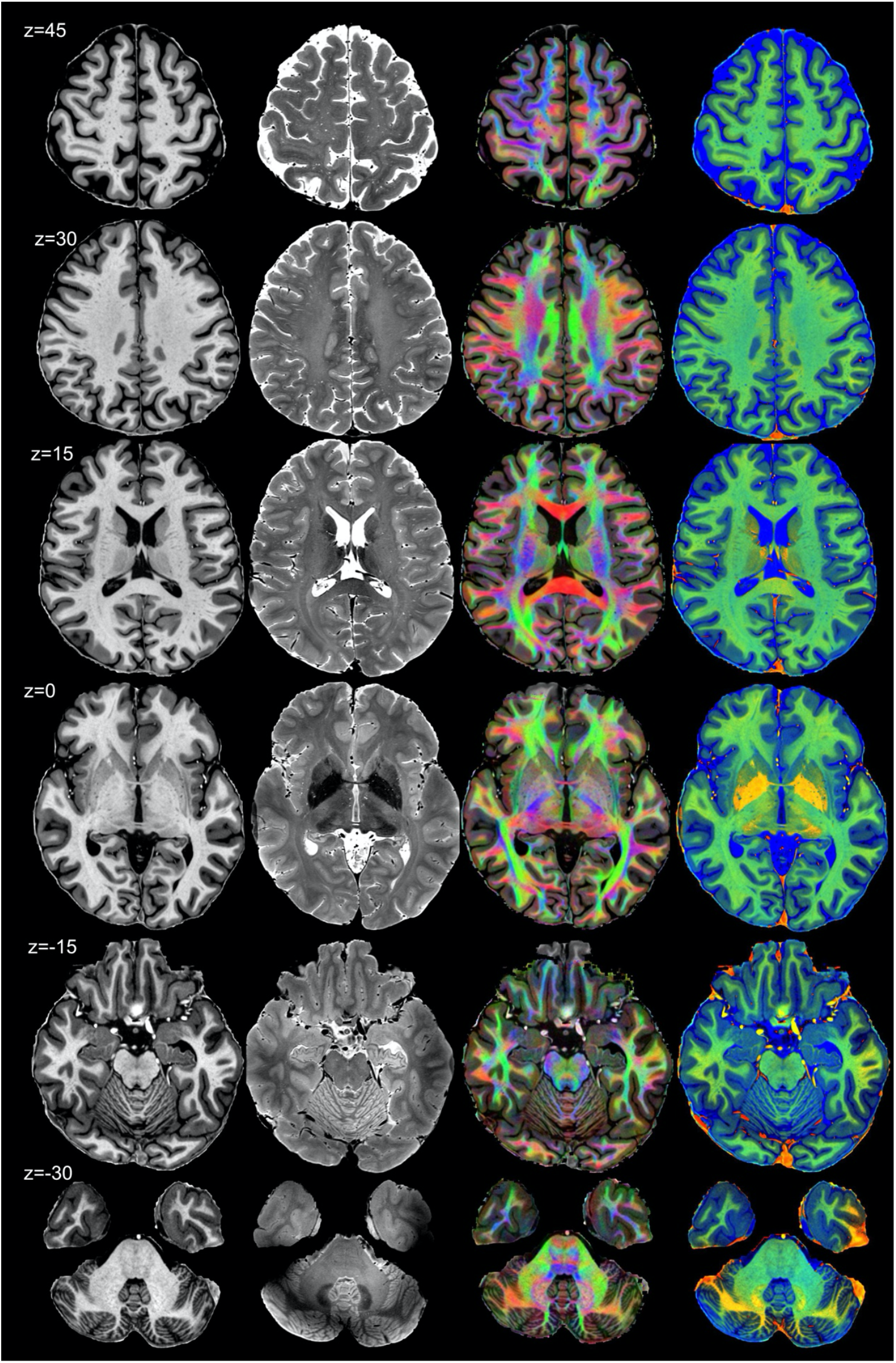
previous page: Horizontal sections from Subject 1. Each column displays a different contrast: T1w, T2w, DEC_T1 and ColorT1T2 and for each row the z position relative to the AC-PC plane is labelled on the left. Note the excellent contrast, structure resolution and sharpness.

**Figure 5.**
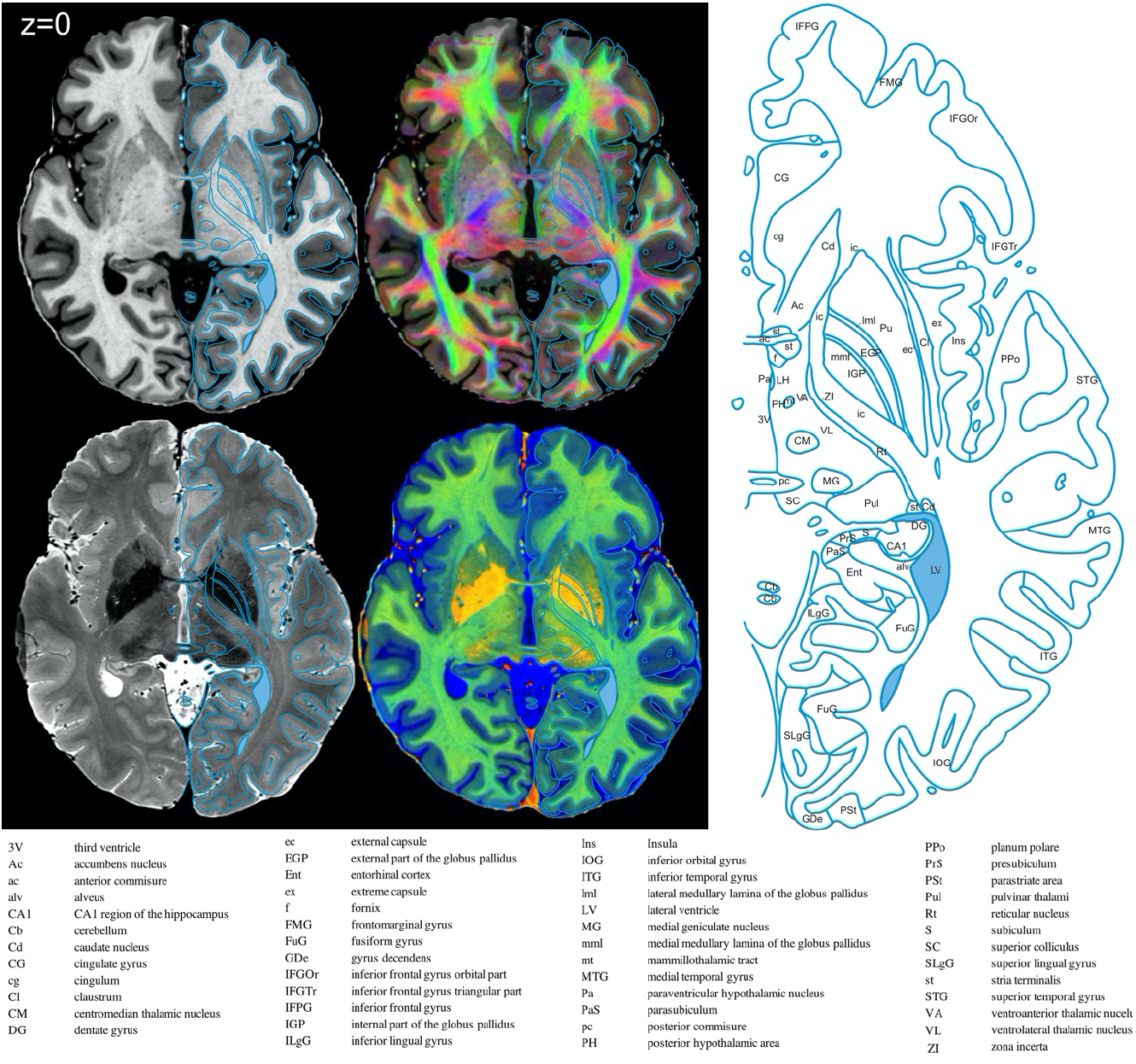
Atlas example from an axial (horizontal) section through the AC-PC line. The delineations identify 53 structures, but we would like to point out that the MRI supports the confident identification of more structures.

Finally, we asked how such MRI delineations compare to delineations based on histology slices that are used for Mai, Majtanik and Paxinos (2016). Figure 6 shows a comparison at the level of the anterior commissure, demonstrating a level of delineations at least equal to the gold standard documents.

**Figure 6.**
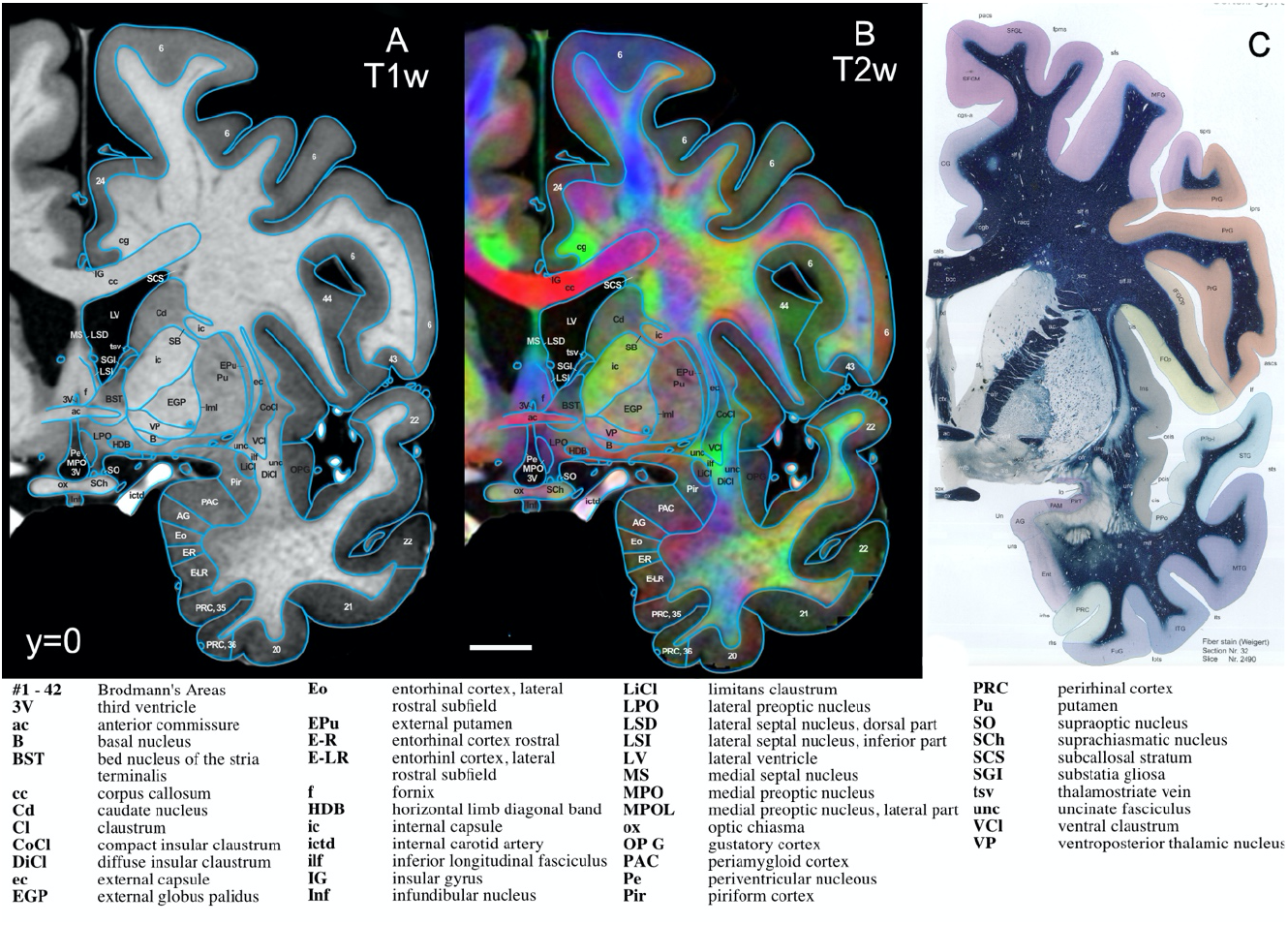
Coronal slices at the level of the anterior commissure (y=0). Panel A shows the T1w contrast and Panel B the DEC_T1 contrast, both with delineations overlayed. Panel C shows the corresponding section from Mai et al (2016) as reference for a comprehensive histology derived atlas (C is reproduced with permission and is exempt from Creative Commons Licence).

To demonstrate justification for these delineations, Figure 7 shows a zoomed in section of the sub-cortex of Figure 6, and we will discuss ten examples. Each one, demonstrates a link between MRI and histology, anatomy, dissections, and/or function, presented as a sample of the scope of referencing material used throughout the rest of the atlas. Figure 7 only shows the facilitating power of the DEC_T1 contrast, albeit the other contrasts are also used in the justifications and are shown in Supl. Figs S3, S4 and S5). The **1**) *internal carotid artery (ictd) is* best delineated by its bright white appearance in T1w and near black appearance in T2w. (Huk & Gademann, 1984). Blood vessels, depending on flow, do not always appear with the same signature in T1 and T2. Blood vessels always appear as vivid red in the ColorT1T2 contrast, hence a good mnemonic and a good distinction from nerves. The **2)** *cingulum bundle (cg)* is identified in Fig 7 by the vivid green colour, signifying an anterior-posterior direction of fibres in the DEC_T1; it also stands out as a darker grey in the T2w. The **3)** *corpus callosum (cc),* like the cingulum, is best identified by its fibre direction, predominantly appearing as vivid red, identifying the mediolateral direction of fibres, with brushes of blue laterally. Modern decisional literature also is used to assist in its delineation (Shah et al., 2021). Leading on from the corpus callosum, the **4)** *superior longitudinal fasciculus, dorsal (slf I)* and **5)** *superior longitudinal fasciculus, central (slf II),* are both rather difficult to identify using only histology or T1w and T2w, but in the DEC_T1 contrast, thanks to directional information, the superior longitudinal fasciculus bundles are identifiable. Functional correlates assist in determining their bounds and validate the blue, identifying a dorsal-ventral direction of fibres, into the superior frontal gyrus (slf I) and green, anterior-posterior, into the medial frontal gyrus(slf I I)(Janelle et al., 2022).The **6)** *inferior longitudinal fasciculus (ilf)* is similarly difficult to identify on low resolution FAC contracts and even via myelin stains; however in the DEC_T1 contrast it is seen as a vivid-green. This colour accurately correlates with the *inferior longitudinal fasciculus* anterior-posterior direction of fibres and is further validated by functional studies (Herbet et al., 2018). The *inferior longitudinal fasciculus* is also delimited by its separation from the LiCl and VCl. Similarly, the **7)** *uncinate fasciculus (unc),* also presents as vivid green on the medial portion of its presence at AC=+0 and blue on its lateral. This correlates with its direction of fibres at AC=+0. The *uncinate fasciculus* is distinguished form the *inferior longitudinal fasciculus,* by the subtle shift to a darker grey in T1w in-between (Bhatia et al., 2017). The **8)** *external globus pallidus (EGP)* is recognisable by its signature dark black in T2w (Zhang et al., 2007), but also visualised by its mottled appearance in the DEC_T1, distinguishable from its surrounding structures. The **9)** *compart insular claustrum (CoCl),* is separated from the *ventral claustrum (VCl)* by a colour shift from turquoise-green to lime-green, corroborated by hodological and histology studies of the area (Watson et al., 2017). Finally, the **10)** *fornix (f)* is a structure less consistent in its directionality as it travels from the midline of the brain underneath the corpus callosum to the hypothalamus, here as it is specifically the ‘columns of the fornix’ it is identified by its dorsal-ventral fibres in blue at the midline.

**Figure 7.**
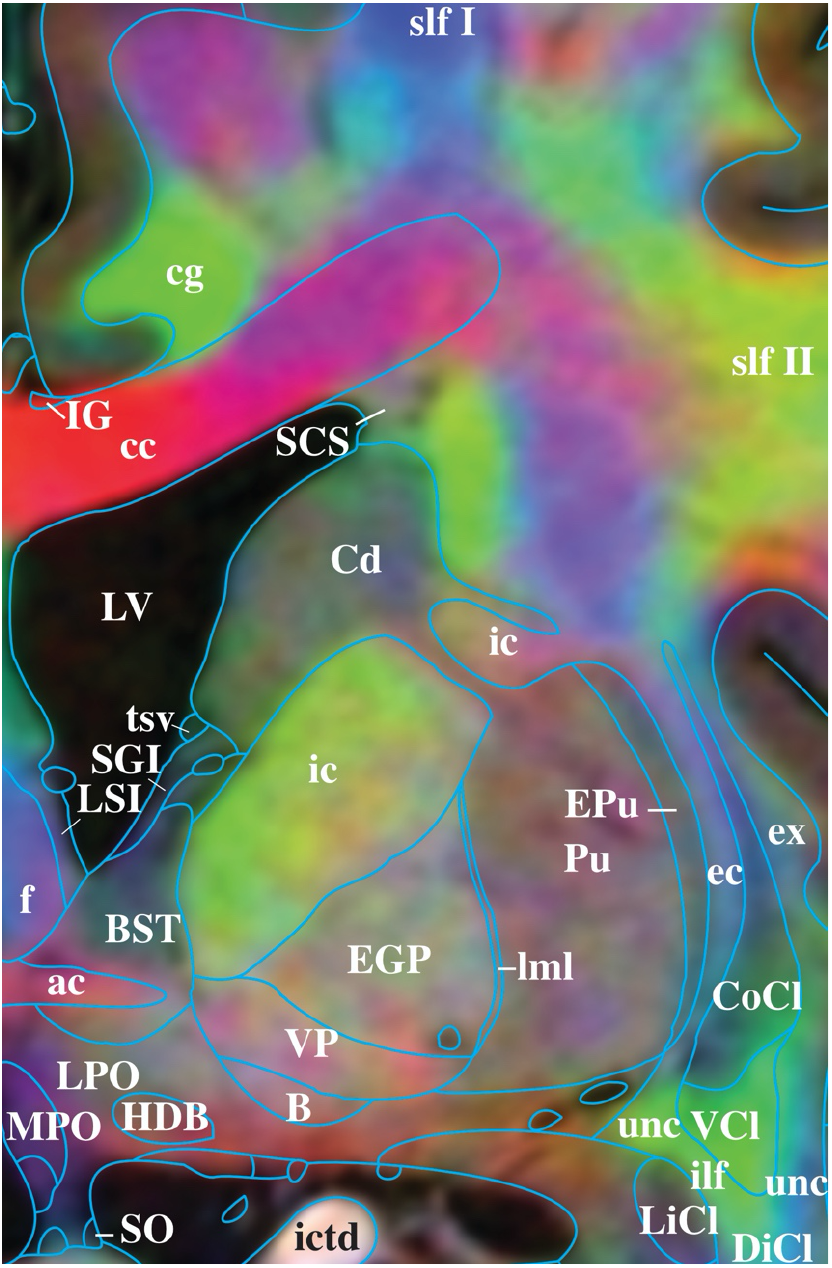
Zoom in on panel B of Figure 6, highlighting the excellent detail and contrast information even at strong magnification, supporting the delineations provided. Legend as in Fig. 6. See also Supplementary Figures S3, S4 and S5 for different contrasts and lower resolutions.

Histological, anatomical, dissectional and functional information, as well as the information provided by the MRI, are used to delineate the striatum and thalamus in Fig 7. It is not purely the MRI contrasts that guide us, but also explicit links to histology. This feature, detail, we argue is what separates HBA from other atlases of the human brain, themselves unique and each contribute different features also used in the disambiguation of gray and white matter structures (Ding et al., 2017; Hawrylycz, Lein, Guillozet-Bongaarts, Shen, Ng, Miller, van de Lagemaat, Smith, Ebbert, Riley, Abajian, Beckmann, Bernard, Bertagnolli, Boe, Cartagena, Chakravarty, Chapin, Chong, Dalley, David Daly, et al., 2012; Sjostedt et al., 2015; Sjostedt et al., 2020; Sunkin et al., 2013b). The final point to note, the key determining these 10 labels, as well as almost every other structure identified throughout the brain, is homology. The human brain and its features are mostly homologous across mammals and to some degree birds. Bird brains having considerable homology to the human, while Primate brains showing virtually no structure unique to one species. With this in mind, the Atlas of the Marmoset (Paxinos et al., 2012), Rhesus (Hartig et al., 2021; Paxinos et al., 2009), Rat (Paxinos & Ashwell, 2018; Paxinos et al., 2021; Paxinos & Watson, 2014; Paxinos et al., 2015), Mouse (Franklin & Paxinos, 2019; Paxinos et al., 2020), and even the Chick (Puelles et al., 2019), are all used to inform homologous structure delineation.

## Discussion

Amongst the most important resources in neuroscience are atlases to navigate the brain. We have acquired magnetic resonance imaging data for the living human of quality that permits detailed delineations. Provided here, are these data (hba.neura.edu.au/data-sets) to serve as the basis for an MRI atlas of the *in vivo* human brain, a dataset with sufficient resolution and contrast to support delineations rivalling histology-based atlases. This is shown in the detailed delineations from this dataset for a coronal and horizontal slice through the anterior commissure, with the DEC_T1 providing additional connectional information that are unavailable in typical histological sections. We argue that HumanBrainAtlas can meet current requirements, offering the identification of structures in a format familiar to researchers and clinicians — *in vivo* MRI.

By averaging multiple low SNR images, we obtained virtually noise free sharp images (Shaw et al., 2019), without the need for additional in scanner hardware. The key lies in sophisticated post-processing, the increased availability of powerful computation hardware and image analysis software, notably the ANTs, FSL and MRtrix packages. Averaging multiple acquisitions, as employed in this project, is a viable approach to improve image quality and resolution. However, it is costly and returns diminish with increasing numbers of acquisitions. Our work prioritised quality, opting for large numbers of repeats; but one could argue this is unfeasible for most applications. However, we suggest it is possible to achieve good enough image quality with significantly less scanning time, and even relying exclusively on 3T. Presumably, acquiring quality MRI will become easier in the next 5 years, for example T1w and T2w at 0.5mm or DWI data at 1mm. In the supplementary materials we provide downsampled examples of Figure 7 (Suppl. Fig S3, S4 and S5), this allows the combination of lower resolution MRI data, yet informed by high resolution segmentation. The images, segmentations, and analysis scripts from the HumanBrainAtlas should facilitate the analysis and interpretation and of such future data.

Our DWI derived images (FAC and DEC_T1) demonstrate that averaging repeated acquisitions in an up-sampled space can provide higher resolution than the native scanning resolution. The detail and quality the FAC images at post averaged resolution of 0.5 is noticeably superior (Fig. 3 bottom left) than the scanning resolution of 1.25 mm (Fig. 3 bottom right). This demonstrates that averaging multiple low resolution images with small misalignments can be used to construct an average image with a resolution higher than that of the acquisition. Presumably the efficacy of this approach rests on the number of images available for averaging. Hence, the super resolution effect was most pronounced in the DWI data, because DWI is an average of multiple images (33) and we averaged multiple DWI acquisitions (10). We suggest a similarly strong effect could be achieved for functional MRI, which is also reliant on many repeated acquisitions, e.g as demonstrated by Bollmann et al. (Bollmann et al., 2017)

Techniques that don’t damage the sample, such as *in vivo* MRI offer advantages over histological atlases, specifically that the images and planes are aligned between contrast modalities, unlike those within histology. In histology, different stained slides are microns apart, at best as little as 20 microns, at worst as much as 1 mm. Each histology slice suffers from sectioning distortions, unique to each slice. Histology, as any other post-mortem technique also suffers from fixation shrinkage and warping. This is unlike the MR images provided here, where each contrast is at every voxel step, with voxel steps analogous to tissue sections in this context, with distortions negligible. Therefore, where histology requires new slices of tissue to compare the cytoarchitecture of the thalamus in different stains, MRI does not. Further, in histology comparing coronal and axial sections, requires a new specimen and MRI does not. These fundamental advantages of MRI were previously insufficient to overcome the resolution disadvantage. Histological preparations still offer significantly higher resolution (< 1micron); while the present resolution MRI is still only 250 microns, this no longer offsets the disadvantages for a whole brain atlas. Instead the advantage of resolution in histology can only be leveraged in very magnified views of small structures, and is most suitable for research investigating neuroanatomy in small subregions of the brain.

Brain atlases are fundamental for neuroscience, being often the template in which other research is placed. Atlases are only contractable and accurate by linking theory from different fields. Readers assume that the delineation were not constructed ignorantly within the atlas they use, but with a comprehensive linking of literature and data. This allows the reader to understand more than purely the anatomy of let’s say the hippocampus, but also its molecular and cognitive relevance, and how it sits within a system—linking a molecular study in the mouse hippocampus region CA1 to a clinical study within the human requires the most accurate and translatable definitions.

We argue that the dataset presented herein and made available for open access satisfies the new needs outlined in the introduction: Enable a high-resolution atlas, free from tissue degradations inevitable in post-mortem material, and in a contrast that is immediately familiar to the user of MRI. The forthcoming atlas, for which we present two sample pages here, will be of value for researchers interested in human or animal nervous systems, clinicians interested in homologies or accurate interventions, and certainly teachers will all be advantaged by this resource. The field is indebted to histology atlases of human brain anatomy have relied on histology (Broadmann, 1909; Büttner-Ennever et al., 2014; Economo & Koskinas, 1925; J. r. Mai et al., 2016; Nieuwenhuys et al., 1978; Talairach & Tournoux), but we argue the future should rely on in vivo MRI.

## Methods

### Subjects

Two male subjects were scanned extensively (up to 20 sessions) for this project. Both were healthy and with no history of neurological or psychiatric conditions. While some scanning parameters differed between subjects, for the most part each subject underwent a similar set of scanning protocols. At the time of scanning, Subject 1 was 45 and Subject 2 was 30 years old.

### Scanning Acquisition Parameters

T1w, T2w and Proton Density (PD) images were acquired on a Siemens 7T MAGNETOM at the Centre for Advanced Imaging using a 32-channel head coil across multiple sessions (up to 12 per subject). Diffusion Weighted Imaging (DWI) data were acquired on a Philips 3T Ingenia TX at the NeuRA Imaging Centre using a 32-channel head coil, again across multiple sessions (up to 10 per subject). More specific details for each protocol are listed below.

### T1w

Three different T1w protocols were used, here called MP2RAGE, Dutch, FLAWS. All three are based on the Siemens WIP 944, a two-inversion MP2RAGE sequence, but using different parameters. The idea is that each sequence will be advantageous for slightly different regions, and the resulting average will benefit from each. The exact scan protocols can be found in the supplementary materials. Briefly the parameters are: a) MP2RAGE Protocol: Voxel size 0.4^3^ mm^3^ isotropic, TR=4300ms, TE=1.8ms, TI1=700ms, TI2=2370ms, FA1=4°, FA2=5°, GRAPPA=2, Echo spacing 5.4ms, Bandwidth 590 Hz/Px, the denoised (UNIDEN) image was used, b) Dutch Protocol: (Fracasso et al. 2016): Voxel size=0.5^3^ mm^3^,TR=6000ms, TE=3.18ms, TI1=1200ms, TI2=4790ms, GRAPPA=3, FA1=8°, FA2=9°, Bandwidth = 630Hz/Px, the PD corrected Inv1 image was used and c) FLAWS Protocol: Voxel size 0.6^3^ mm^3^,TR=5000ms, TE=1.49ms, TI1=620ms, TI2=1450ms, FA1=4°, FA2=8°, GRAPPA=3 partial Fourier 6/8 Bandwidth = 630Hz/Px, the PD corrected Inv2 image was used.

### T2w

T2w images were collected with a 3D TSE sequence (SPACE) using the Siemens WIP692; again, detailed parameters can be found in the supplementary materials. Briefly, TR=1330ms, TE=118ms, GRAPPA 3, SPAIR fat suppression, 384 slices, FOV 256mm, Matrix size 640×640, resolution 0.4^3^ mm^3^, Bandwidth = 521Hz/Px.

### Proton Density (PD)

In each session, one proton density (PD) scan was collected, Voxel size 1 mm^3^, TR=6.0ms, TE=3.0ms, GRAPPA=2. One PD scan was collected during each scanning session in order to correct for intensity inhomogeneities present in our T1w scans (Van de Moortele et al, 2009).

### Diffusion-Weighted Imaging (DWI)

DWI data were acquired on a 3T Phillips Achieva CX, at the NeuRA Imaging Facility in Randwick, Australia using a diffusion tensor imaging (DTI) echo planar imaging (EPI) sequence. Native scan resolution was 1.25 mm^3^ isotropic, field of view (FOV) 240×200mm and 118 slices, 32 directions, 4 b-factor averages, B-val=1000, TE=60ms, TR=26.5s, SENSE=3, SPIR (Spectral Saturation with Inversion Recovery) fat saturation, fat shift direction A to P, distortion corrected using inverse blip scan, resulting in a total scan time of 52 minutes. Ten scans were acquired with one scan per session to ensure maximum subject compliance with minimal motion.

### Data Analysis

### T1w and T2w pre template preprocessing

Each T1w scan type (MP2RAGE, Dutch, Flaws) was preprocessed independently through the following steps:

1. Applying the ImageMath command from the ANTs toolbox to truncate the luminance intensities of each scan with 0 as the lower quantile and 0.999 as the upper quantile.
2. Using the robustfov script from FSL to reduce file size by removing unnecessary parts of the scan (neck, nose, etc).
3. Upsampling (b-spline interpolation) the voxel size to 0.25 mm^3^ to ensure voxel sizes were uniform across different scan protocols and modalities, this decreases the effect of blurring caused by ‘reslicing’ or ‘resampling’ and allows some degree of super resolution by integrating information over multiple frames (Farsiu et al., 2004; Manjón et al., 2010; Tsai, 1984; Van Reeth et al., 2012) to increase detail.
4. Skull stripping was undertaken to improve alignment by removing parts of each scan that did not include cortex. We also conducted skull stripping due to the large file size of our raw scans (~1-2Gb), as skull stripping decreased file sizes considerably (~300Mb). For this we used HD-BET to create a brain mask for each scan (Isensee et al., 2019). To avoid this mask removing brain areas with low signal (e.g., temporal cortex), we also created a skull stripped variant with the brain mask inflated by 30mm. Accurate skull stripping is critical, and results were carefully inspected. When the skull strip was inaccurate for a specific scan, an accurate skull strip mask from another dataset was used. For this the two datasets were aligned and when this alignment was successful the accurate mask was transformed as well. Also inaccuracies in skull stripping were manually corrected using the ITKSnap software.
5. Ensuring all the dimensions of each scan was 1024. As in Step 3, this was used to ensure uniform dimensions across different scan protocols and modalities. With background being zero values, this did not increase file sizes as the files were saved in a zipped format (nii.gz).
6. Proton Density Correction. We used the method proposed by Van de Moortele et al (Van de Moortele et al., 2009) to correct for luminance inhomogeneities in our T1w images. Proton Density images were collected during the same scan session for each T1w image, and the T1w images acquired using the ‘Dutch’ and the ‘FLAWS’ protocol were divided by the aligned Proton Density image. This yielded images with much reduced inhomogeneities, and high grey/white contrast.
7. All T1w and all T2w images were then aligned using FLIRT and a linear average was generated using the fslmerge command with the –t flag. This created one unbiased linear average for each for T1w and T2w set of scans, avoiding influence by the order in which the scans were collected. This linear average was used as a starting point for symmetric groupwise template generation.

### Template generation

Symmetric group-wise normalisation was conducted using Advanced Normalization Tools (ANTs), specifically using the antsMultivariateTemplateConstruction.sh script. The parameters used to align each scan included a Cross Correlation similarity metric a Greedy SyN transformation model used for non-linear registration, 20×15×5 was the maximum number of steps in each registration, the gradient was 0.1, and the total number of iterations was 2. These settings were based of the settings used by (Lüsebrink et al., 2017). Again, each modality (T1w, T2w, DWI) was processed separately and aligned afterwards using ANTs non-linear registration. A non-linear alignment was chosen after the results of affine alignments were found to be good, but still marginally suboptimal. This was specifically noticeable in the computed ColorT1T2 which we observed to be sharper after non-linear alignment.

### Diffusion-Weighted Imaging (DWI)

DWI data was analyzed using MRtrix 3.2 (Tournier et al., 2019) and ANTs. Firstly, each scan was upsampled by a factor of 2×2 in the inplane direction (to 0.625 mm × 0.625 mm × 1.25 mm) and then preprocessed using the dwifslpreproc script providing top-up distortion correction and eddy_cuda correction (Jenkinson et al., 2012). Then the preprocessed data was upscaled across slices, again by a factor of 2, so the final output had an isotropic resolution of 0.6123^3^ mm^3^. This upsampling strategy was chosen to increase the spatial detail through averaging repeated acquisitions. Doubling the across slice resolution is incompatible with eddy correction, hence only the inplane direction can be upsampled before eddy correction. For each of the 10 upsampled preprocessed DWI datasets, a mean DWI image, a FAC (Fractional Anisotropy Color) image as well as a FOD (Fibre Orientation Distribution, dwi2fod) using dhollander algorithm (Tournier et al., 2019)) was calculated.

Subsequently, the 10 mean DWI images were aligned using the ANTS MultivariateTemplateConstruction.sh script, resulting in high-resolution high-quality mean DWI essentially displaying a T2*w contrast with good image contrast and detail. The transforms estimated from this were then applied to the 10 FAC images (via 10 ID images) and the 10 FOD images using mrtransform function from MRtrix which ensured that the FOD vectors were transformed correctly (using the option -reorient_fod yes). FAC images and FOD were then averaged in the DWI template space using mrcalc. This intermediate space was used to save RAM and compute resources as averaging 10 FOD images at 0.25mm required above 128GB of RAM. Then, all DWI data were transformed into the final ACPC template space (0.25 mm) using ANTs for the FAC and mean image and ID files and MRtrix for the FOD. Finally, from the average FOD a DEC (direction encoded color) image was calculated using the T1w template for panchromatic sharpening (fod2dec -contrast) which also transferred the DEC_T1 image voxel matching into template space.

### Template alignments, multi contrast images, final contrasts

Great care was taken to achieve optimal alignment of these different image modalities, overcoming the minute distortions of each. We computed multi-contrast images, to validate the accuracy of these alignments, where alignment inaccuracies would introduce blur into these multi contrast images. Firstly, the image “ColorT1T2” was calculated from the T1w and T2w by manipulating the RGB (red, green, blue) channels of the image. The red channel of the image was calculated as T1w/T2w (using Matlab). This was thresholded at the 96^th^ percentile, as the division resulted in some extremely large values in the background. The green channel of the image is the T1w, and the blue channel the T2w image. This colour scheme was chosen because it highlights some blood vessels in red greatly simplifying the discrimination of blood vessels and nerve fibres in the manual segmentation.

The second multi-contrast image is called DEC_T1 and combines the T1w and the DWI data. Specifically, it is the output of the fod2dec function encoded image derived from the FOD (Dhollander et al., 2015).

**Table 1.** Summary of the MRI scans included in the HumanBrainAtlas project.

| name | derived from scans | description |
| --- | --- | --- |
| T1w | 7T MP2RAGE, DUTCH, FLAWS | T1 weighted scan. The main reference contrast, derived by averaging all T1w scans of sufficient quality. |
| T2w | 7T 3D TSE sequence (SPACE) | T2 weighted scans. |
| DWI_average | 3T SPIR EPI | Average of the distortion-corrected EPI images. Mostly used for aligning the DWI data to the T2w (and hence the T1w) image. |
| FAC | 3T SPIR EPI | For every DWI scan a FAC image was calculated. These 10 FACs were aligned with the T2w and averaged. |
| DEC_T1 | T1w, 3T SPIR EPI | Direction encoded fractional anisotropy reconstructed from the FOD (fod2dec -contrast). |
| ColorT1T2 | T1w and T2w | A derived image combining the T1w and T2w contrast — helpful for segmentation and highlighting of blood vessels. |

### Detailed documentation of the processing scripts

We provide a detailed “How to” like description of the data processing online:

1. WITHOUT an input template: https://hackmd.io/DQbUc5lJQ_u9qCyd3Q3kSA
2. WITH an input template: https://hackmd.io/qKZtbfpSTnGzSebiQW3KfQ
3. Using SLURM to run ANTs: https://hackmd.io/b5YqltBPRgqxXl83ZVbjcQ
4. DWI data: https://hackmd.io/0W2D1db_QROG_HU9Vv4-JA

### Delineation of neuroanatomy

For each 0.25 mm^3^ voxel a set of three orthogonal planes were extracted: coronal, sagittal, and axial (horizontal). For each plane, a series of images were compiled per voxel step, from each of their respected ranges, i.e., for coronal, from 65 mm before the anterior commissure (−65AC) to 80mm after the anterior commissure (+80AC). These MR slice images are analogous to histological sections allowing direct and practical comparisons with existing delineated histology slices.

With these series of images, across all contrasts a set of four ‘virtual’ fiduciary marks were placed in the four corners of the image, these ensure that as we delineate through the series, we are always on the same side, aspect ratio, and alignment. The T1w contrast of each voxel step is overlaid with 0.05 mm drafting film (Flat and Rotary Co., Ltd.) to permit superimposition of each other contrast, where in conference with co-authors, the signatures of structures are identified and then drawn.

Once satisfied with each voxel step/slice, we move to the next, and repeat the process until the range is complete and then repeat it again for each orthogonal plane. Albeit, this is not done in complete serialisation, as we have determined following a structure, or small set of structures, from start to finish, across orthogonal planes is most efficient and accurate. These tracings are then digitised using Adobe software (Adobe Inc., 2019 Adobe Suite). At this digitisation step, a final sweep through the now diagrams is done to harmonize delineations from level to level.

## Rights and permissions

This work is licensed under the Attribution-NonCommercial-ShareAlike 4.0 International License. The images and material in this article are included in the article’s Creative Commons license, unless indicated otherwise in the credit line; if the material is not included under the Creative Commons license, users will need to obtain permission from the license holder to reproduce the material. To view a copy of this license, visit https://creativecommons.org/licenses/by-nc-sa/4.0/.

## Supplementary Materials

**Table S1.** Sequence parameter summary for Subject 1.

|  |  | Voxel Size | Dimensions | Usable Scans/Number of Scans Collected | TR | TE | FOV |
| --- | --- | --- | --- | --- | --- | --- | --- |
| T1w | Dutch | 0.50 x 0.50 x 0.50 | 312 x 468 x 500 | 6/6 | 4.3 | 0.00183<br>0.01 |  |
|  | Flaws | Scans 1 to 3:<br>0.530 x 0.525 x 0.525<br>Scans 4 to 6:<br>0.570 x 0.5721 x 0.5721 | Scans 1 to 3:<br>256 x 374 x 400<br>Scans 4 to 6:<br>268 x 376 x 402 | 3/6<br>(scans 1-3 were discarded due to folding) |  |  |  |
|  | MP2RAGE | 0.400 x 0.4015 x 0.4015 | 396 x 528 x 528 | 12/12 |  |  |  |
| T2 |  | Scans 1 to 4:<br>0.33 x 0.3281 x 0.3281 | Scans 1 to 4:<br>384 x 640 x 640 | 8/12 |  |  |  |
|  |  | Scans 5 to 8:<br>0.4 x 0.4 x 0.4 | Scans 5 to 8:<br>384 x 640 x 640 | (scans 1 to 4, FOV to small) |  |  |  |
| DWI |  | 1.25 x 1.25 x 1.25 | Scan 1,3-11:<br>192x192x118x33<br>Scan 2 192x192x36x33 | 10/11<br>Scan 2 was partial coverage aiming to optimize distortion but that failed. | 26.5s | 60ms | 240x240x147.5mm |

**Table S2.** Sequence parameter summary for Subject 2.

|  |  | Voxel Size | Dimensions | Usable Scans/Number of Scans Collected | TR | TE | FOV |
| --- | --- | --- | --- | --- | --- | --- | --- |
| T1w | Dutch | 0.50 x 0.50 x 0.50 | Scans 1 to 2:<br>320 x 468 x 500<br>Scans 3 to 5:<br>312 x 468 x 500 | 5/5 | 4.3 | 0.00183<br>0.01 |  |
|  | Flaws | Scans 1 to 4:<br>0.64 x 0.64 x 0.64<br>Scans 5 to 8:<br>0.570 x 0.5721 x 0.5721 | Scans 1 to 3:<br>256 x 374 x 400<br>Scans 4 to 6:<br>268 x 376 x 402 | 7/8<br>(scan 7 - file corrupted) |  |  |  |
|  | MP2RAGE | 0.400 x 0.4015 x 0.4015 | 396 x 528 x 528 | 12/12 |  |  |  |
| T2 |  | Scans 1 to 4:<br>0.5 x 0.5 x 0.5<br>Scans 5 to 11:<br>0.4 x 0.4 x 0.4 | Scans 1 to 4:<br>320 x 512 x 512<br>Scans 5 to 8:<br>384 x 640 x 640 | 8/12<br>(scans 1 to 4, FOV too small) |  |  |  |

**Fig. S1.**
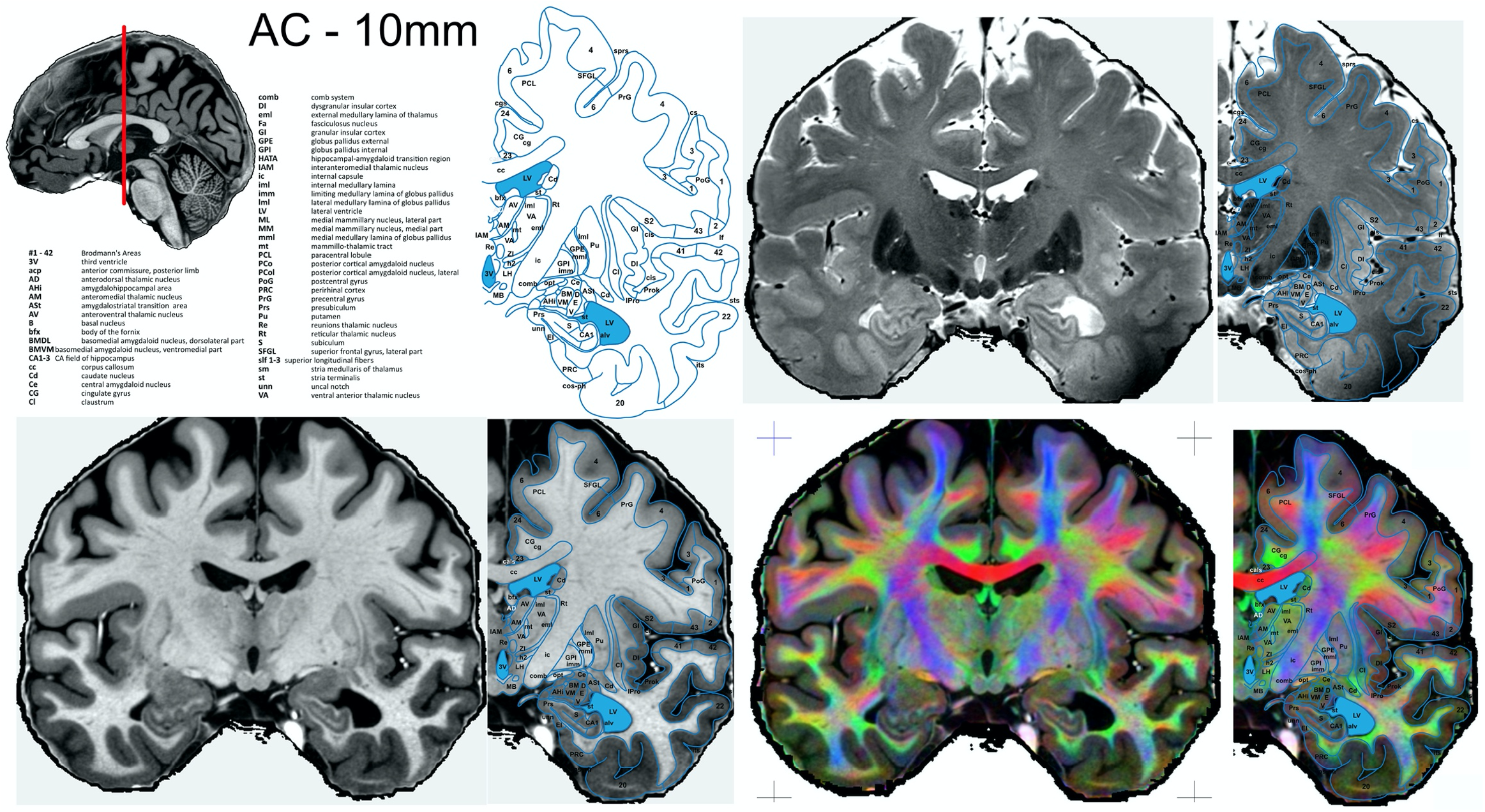
Example atlas page at position AC – 10 mm, that is 10 mm posterior of the anterior commissure or y = -10.

**Fig. S2.**
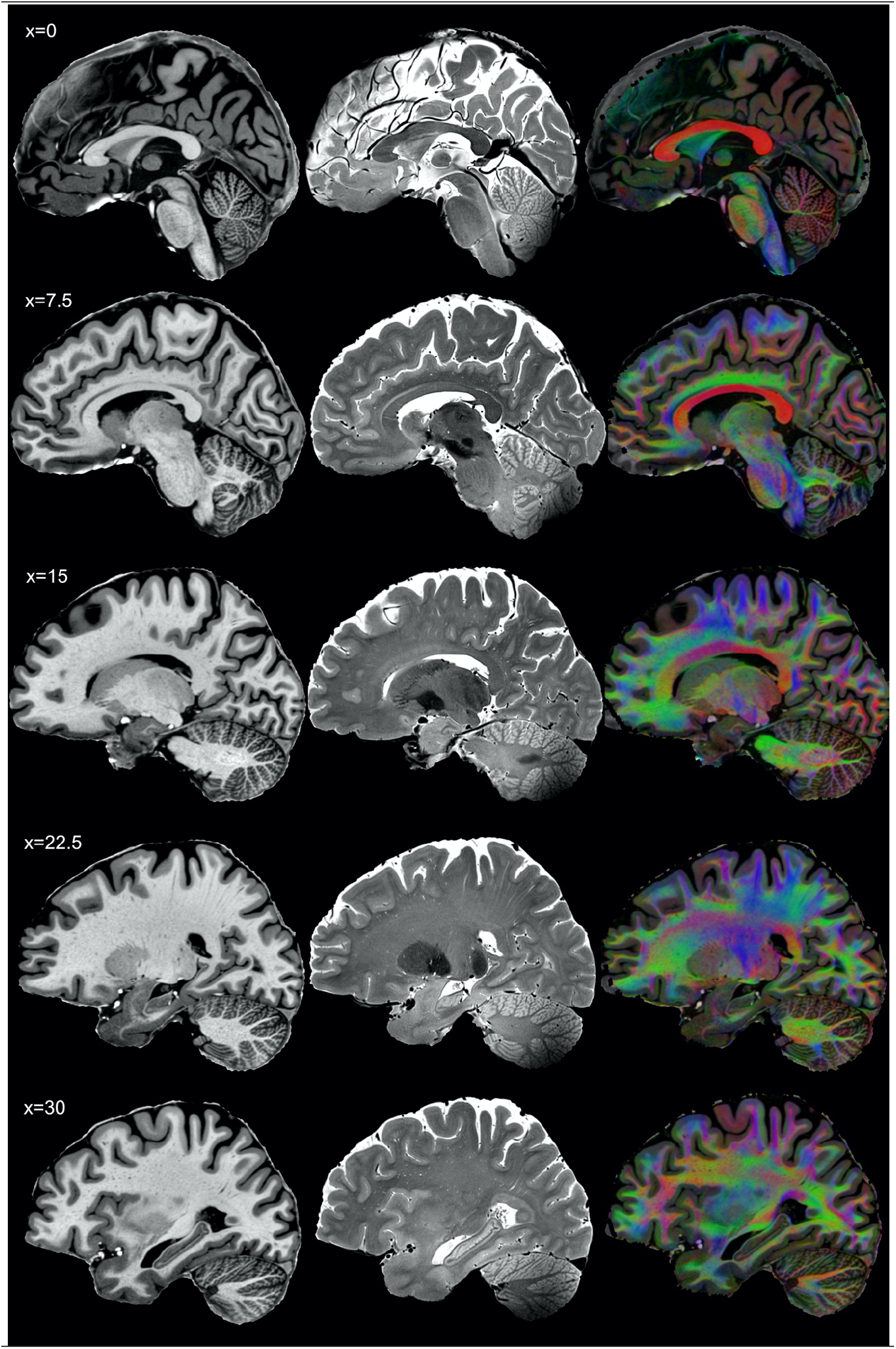
Series of sagittal slices, complementing the horizontal and coronal sections presented in the main manuscript.

**Fig. S3.**
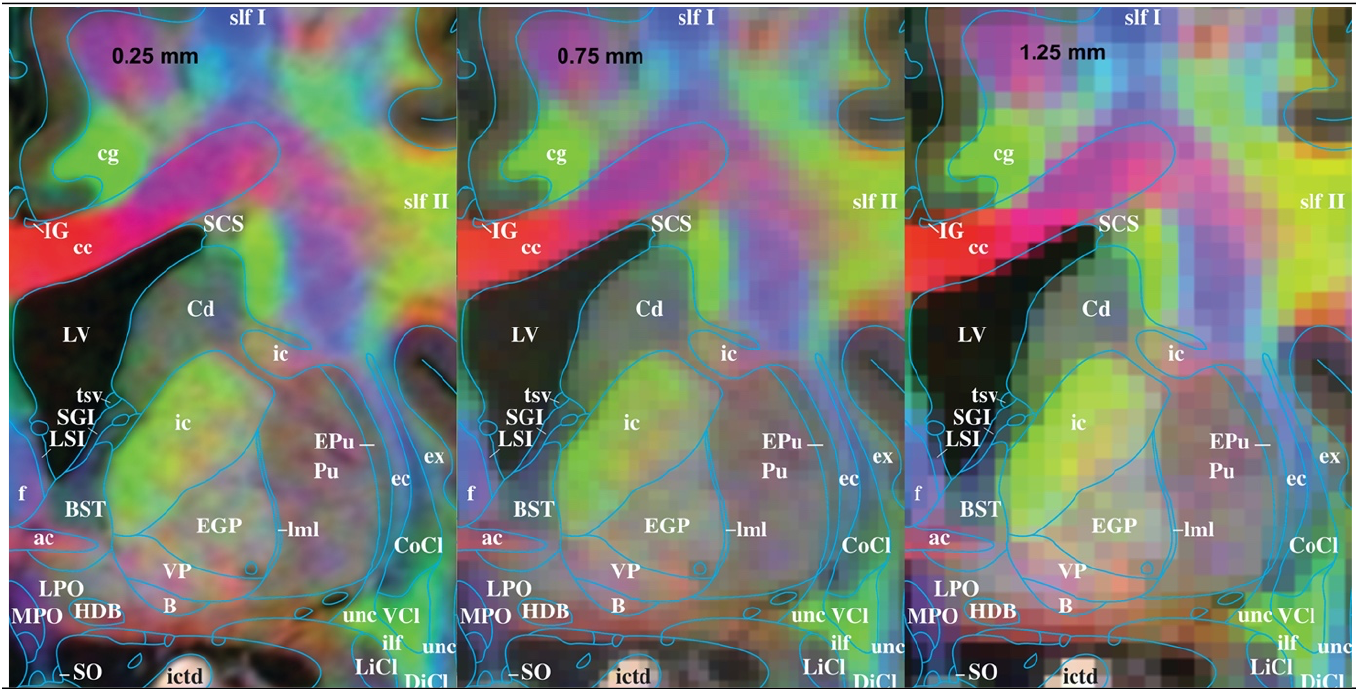
Comparing DEC_T1 images at 0.25 mm, 0.75 mm and 1.25 mm respectively. Despite clear loss of detail, the there is still significant structural detail at 0.75 and even 1.25mm. The interpretation of lower resolution images is significantly aided by the high-resolution example we provide. Our high-resolution data identifies the anatomical signatures, that allows the localisation of structures such as the bed nucleolus of the stria terminalis (BST), the internal capsule (ic), the external globus palidus (EGP) and the pulvinar (Pu) which all show different fibre orientation profiles even in the low resolution case.

**Fig. S4.**
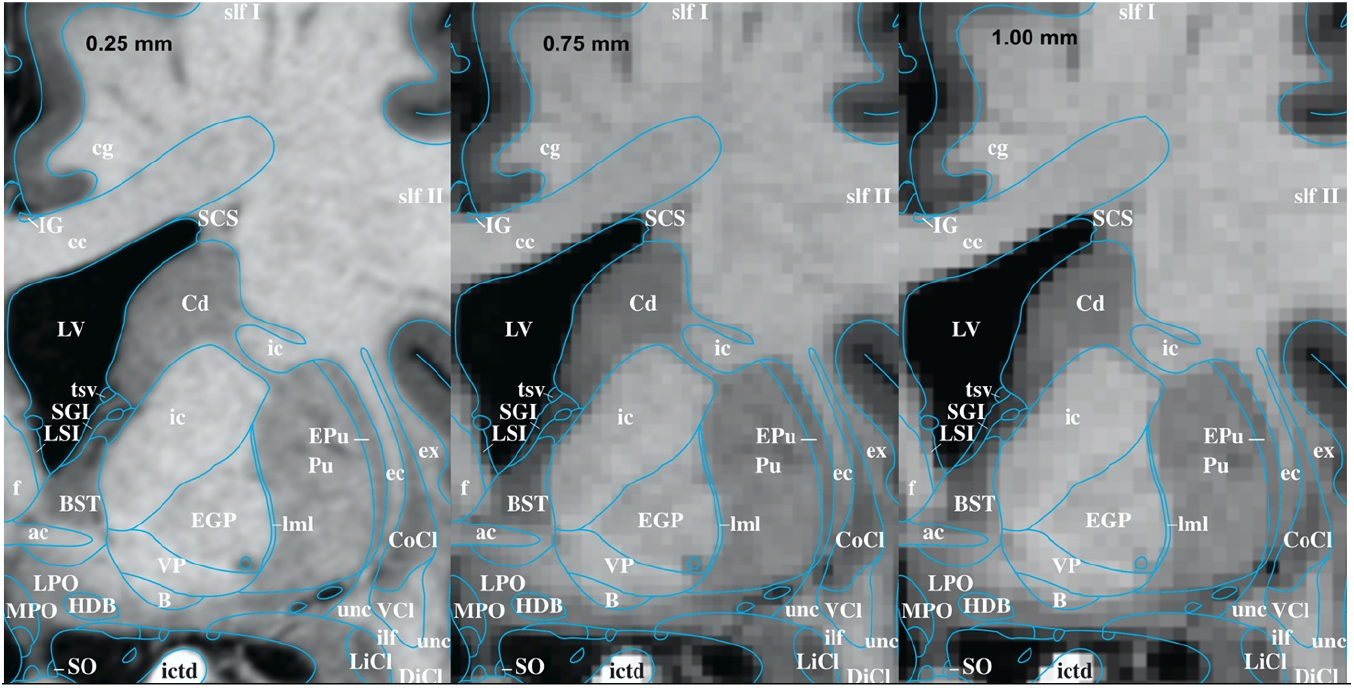
Comparing T1w at 0.25 and 0.75 and 1 mm. These resolutions were chosen as 0.75 and 1mm are rather commonly chosen for high quality T1w images.

**Fig. S5.**
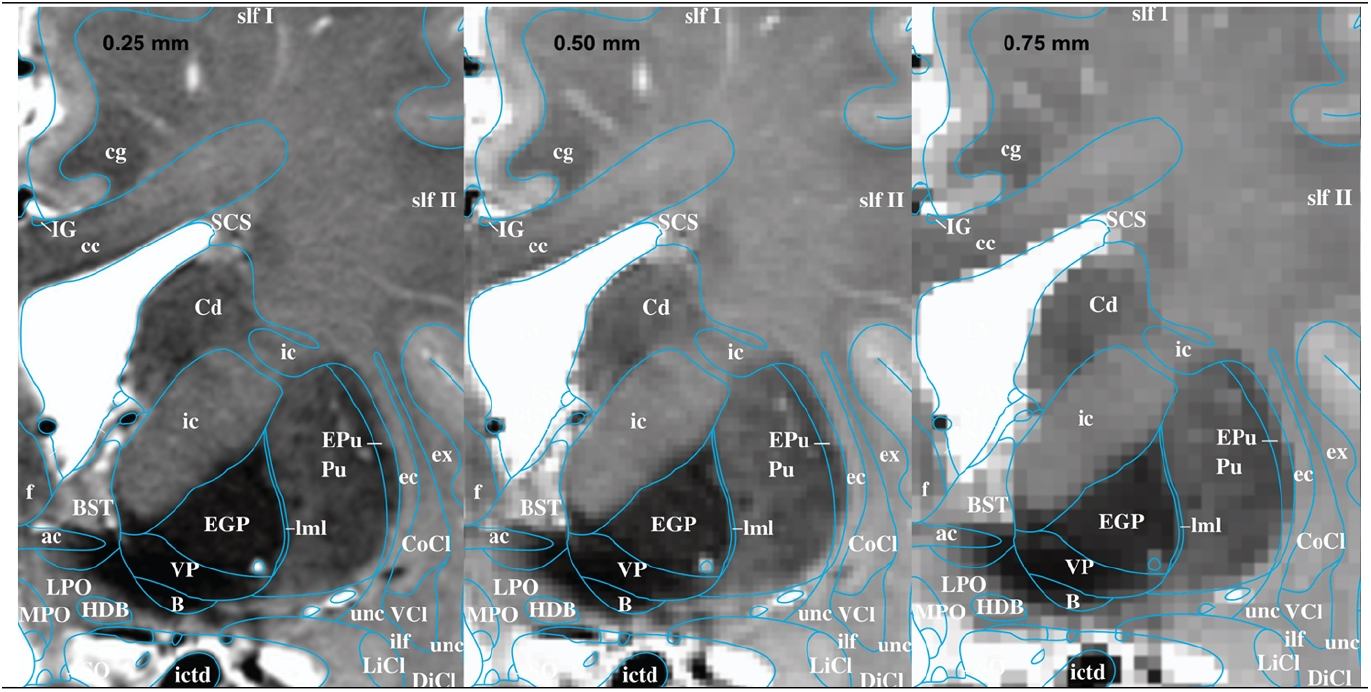
Comparing T2w resolutions, here 0.25mm, 0.5mm and 0.75mm. Illustrating the loss of detail in T2w images. The high resolutions were chosen since such high resolutions are more practical for T2w than for T1w imaging sequences.

